# Autistic kinematics diverge from the power laws that typically govern movement

**DOI:** 10.1101/2023.03.23.532745

**Authors:** Jennifer L. Cook, Dagmar S. Fraser, Lydia J. Hickman, Rebecca Brewer, Dongsung Huh

## Abstract

Extant work reliably demonstrates that autistic individuals move with increased jerk (where jerk concerns change in acceleration). Although it follows that autistic movement may therefore diverge from fundamental power laws that govern movement, this hypothesis has not been directly tested to date. This lack of insight holds back progress in understanding the mechanisms underpinning differences in autism in motor control particularly with respect to movement jerkiness. Here we investigated whether movements executed by autistic adults diverged from the typical power law relationship that links movement speed and curvature. x and y position of the stylus tip was recorded at 133 Hz while 21 autistic and 19 non-autistic age-, intelligence quotient- and sex-matched adults traced, on a tablet device, a range of shapes that varied in angular frequency from 2/33 (spiral-like shapes) to 4 (square-like shapes). The gradient of the relationship between speed and curvature for each angular frequency-defined shape is reliably predicted by a set of mathematical equations often referred to as fundamental power laws thus, to assess deviations from power laws, we compared autistic and non-autistic participants in terms of speed-curvature gradients. To gain insight into potential mechanisms underpinning any differences we also used fast Fourier transform to explore amplitude spectral density across all angular frequencies. Compared to non-autistic adults, autistic adults exhibited significantly steeper speed-curvature gradients. Fast Fourier transform further revealed that non-autistic participants exhibited highly precise modulation of speed oscillations around the target frequency. For example, when drawing an ellipse their speed profile was dominated by speed changes in a band centred around the angular frequency 2 with minimal changes in other bands. Autistic adults, in contrast, exhibited less precise modulation of speed oscillations around the target frequency, a result that is reminiscent of a literature reporting broader auditory filters in autistic individuals. These results evidence, in autistic adults, a deviation from the power laws that typically govern movement and suggest differences in motor cortical control policies and/or biomechanical constraints.

## Introduction

Research into autism spectrum disorder (referred to as autism^[1]^ hereafter)^1,2^ has classically focused on social cognition, language and communication. A burgeoning field, however, documents bodily movement and motor control differences.^3–6^ These motor differences include functional challenges with both ‘fine’ motor tasks such as handwriting,^7–10^ and ‘gross’ motor tasks such as riding a bike or throwing a ball.^11^ Motor control difficulties in autism can also contribute to social communication challenges, including poor interpretation of autistic movements by neurotypical individuals.^5,12–14^ Motor control difficulties may therefore comprise a dual barrier that prevents autistic people from achieving optimal social, educational and health outcomes and makes it more difficult for non-autistic people to help.

Studies of motor control suggest that complex movements, once decomposed, consist of common signatures; thus, by studying simple movements it is possible to gain insight into more complex action sequences.^15^ Our previous work revealed differences in the kinematics of arm movements in autism.^16^ We found that, for simple movements from one point in space to another, non-autistic individuals accelerate and decelerate gradually, producing smooth, minimum-jerk, kinematic profiles,^17,18^ by contrast autistic individuals accelerate and decelerate rapidly, producing jerky movements. We subsequently replicated this result in an independent sample,^14^ and other research teams have reported atypically jerky arm,^19,20^ leg^21^ and head^22^ movements in autistic adults and/or children. Extant work therefore reliably demonstrates that, compared to non-autistic individuals, autistic individuals move with increased jerk.

The finding that autistic individuals move with increased jerk is striking because it potentially indicates a departure from highly replicable laws of motion which govern human and non-human movement. One of these laws, referred to as the one third power law^[2] 23^ describes the relationship between movement speed and trajectory curvature. When drawing an ellipse, for example, people move slowly around the “corners” compared to straight segments.^24–29^ The one third power law states that there is a negative relationship between (log) curvature and (log) speed (**Fig. 1**): as curvature increases, speed decreases. The gradient of this negative relationship is remarkably reliably (negatively) predicted by the ‘angular frequency’ ^[3] 25,30^ of the shape of the movement trajectory.^25^ That is, the gradient of the speed-curvature relationship is steep for shapes, like spirals, with low angular frequency and shallow for shapes, like rounded squares, with high angular frequency (**Fig. 1**). Since these mathematical laws have been observed for hand drawing, pursuit eye movements, speech and walking,^23,31–33^ and have been demonstrated across species including humans, non-human primates,^25,31,34–36^ *drosophila* larvae^29^ and the bumblebee *Bombus terrestris audax*,^37^ it seems that they comprise fundamental, cross-species laws of movement.

**Figure 1.**
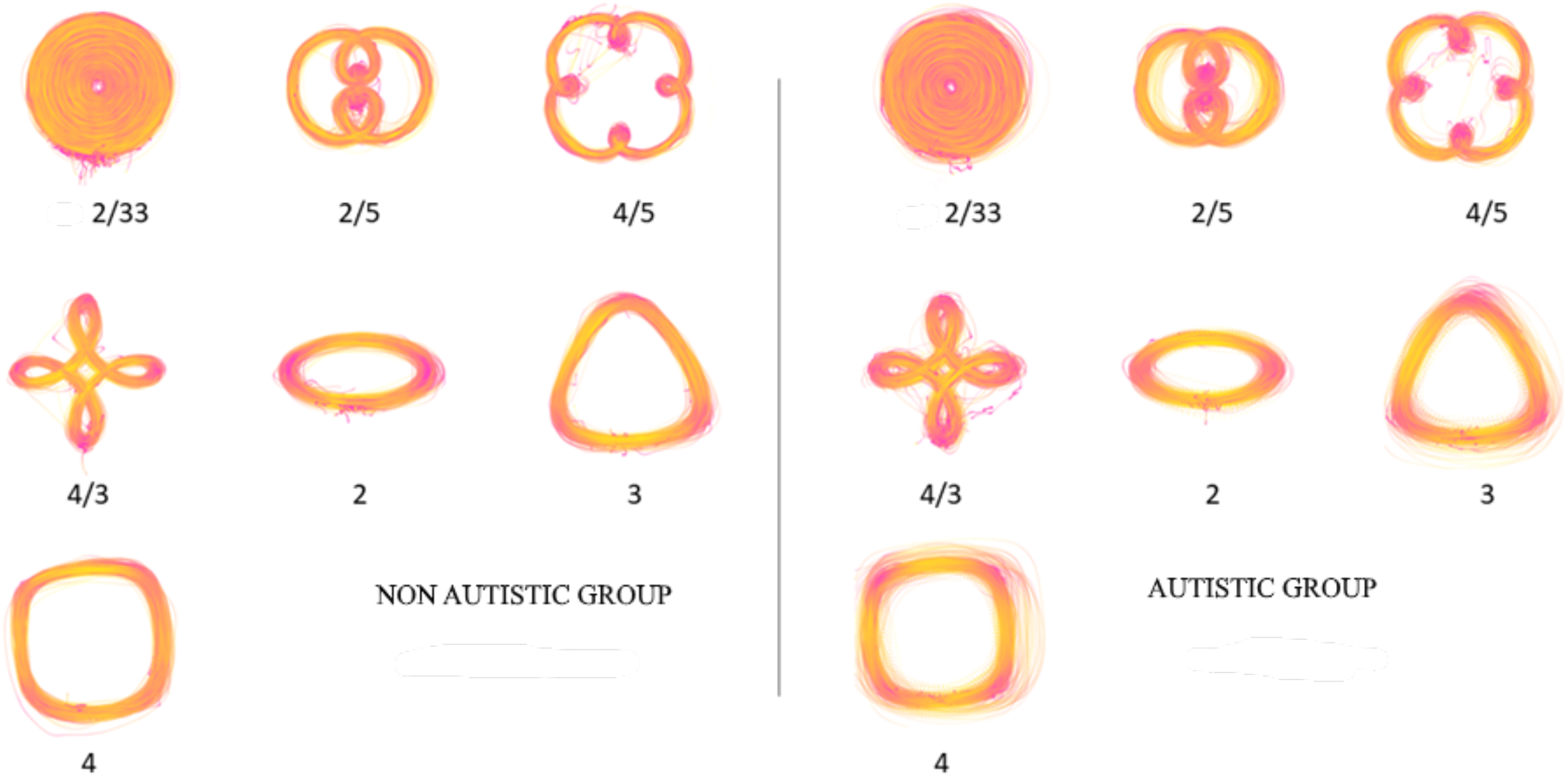
Raw Movement Trajectory Data. Raw trajectory data for all participants in the non-autistic (*left*) and autistic (*right*) groups, for all trials. The colour scheme represents speed from dark pink (low speed) to yellow (high speed). Note that more tightly curved trajectories tend to be represented as dark pink (i.e., low speed).

Since autistic movements tend to be atypically jerky (where jerk concerns change in acceleration) it is possible that they diverge from the power laws that typically govern movement. Studies to date have not, however, directly tested this hypothesis. Existing studies have not calculated speed-curvature gradients for autistic movements. Furthermore, extant studies of autistic movement have focused on shapes with an angular frequency of two (point-to-point movements and ellipses) and have therefore not been able to assess whether autistic movement diverges from the typical power law relationship across the full angular frequency spectrum. Importantly, jerky movements do not *necessarily* result from an atypical power law relationship between speed and curvature. Jerky movements may also be a consequence of decomposing shapes into a greater number of submovements^38–40^ and/or from moving atypically quickly or atypically slowly.^41,42^ Since the speed of movement and the number of submovements are theoretically independent of speed-curvature gradient, it may be that although autistic individuals exhibit atypically jerky kinematics, this is not indicative of a violation of the typical power laws that govern movement.

Understanding whether autistic movements deviate from fundamental power laws has important implications. Deviations from classic power laws may indicate differences in motor cortical control policies^17,43^ and/or biomechanical constraints.^34^ In contrast, simply moving at an abnormal speed, in the absence of violations of power laws, may point to other mechanisms such as an atypical subjective cost function whereby moving fast is particularly expensive/cheap.^40^ Likewise, decomposing movements into a greater number of submovements (without violating power laws) could be a consequence of differences in perceptual processing styles.^44^ To progress towards an understanding of the mechanisms underpinning motor control in autism, we must assess whether movements deviate from the fundamental power law relationship.

Here we investigated whether movements executed by autistic adults diverged from the typical power law relationship linking movement speed and curvature. We recorded x and y position at 133 Hz while autistic and non-autistic participants traced, on a tablet device, a range of shapes that varied in angular frequency from 2/33 (spiral-like shapes) to 4 (square-like shapes). We calculated the gradient of the relationship between speed and curvature for each angular frequency-defined shape and compared gradients between age-, intelligence quotient (IQ)- and sex-matched autistic and non-autistic adults. To gain insight into potential mechanisms underpinning any differences we also used fast Fourier transform to explore amplitude spectral density across all angular frequencies.

To précis our results, we observed that, compared to non-autistic participants, the gradient of the relationship between speed and curvature was significantly steeper for autistic individuals. Fast Fourier transform further revealed that autistic participants exhibited reduced precision of speed oscillations around the target frequency. These results evidence, in autistic individuals, a deviation from the power laws that typically govern movement.

## Materials and Methods

### Participants

Data were collected from 21 participants with a clinical diagnosis of autism and 19 non-autistic participants. The sample size was based on a previous result in which we observed a significant difference in jerk (t(27) = 3.28, P = 0.003, Cohen’s d = 1.26) with 15 non-autistic participants and 14 autistic participants (Cook et al., 2013).^16^ Using GLIMMPSE^45^ we calculated that 6 participants are necessary in each group to have 99% power to detect an equivalent effect. To guard against the “winners curse”^46^ we planned to recruit at least triple the recommended sample size.

All participants in the autistic group had a clinical diagnosis from an independent clinician. In addition, participants in the autistic group completed the Autism Diagnostic Observation Schedule-2^47,48^ with a trained administrator and reached the criteria for autism or autism spectrum. Participant groups were matched on full scale Intelligence Quotient (IQ), as measured by the Wechsler Adult Intelligence Scale^49^ for the autistic group and the Wechsler Abbreviated Scale of Intelligence^50^ for the non-autistic group. There were no significant differences between the groups in terms of age, IQ or sex. The autistic group had significantly higher Autism Quotient (AQ)^51^ scores than the non-autistic group (see **Table 1**). All experimental procedures were approved by the University of Birmingham Research Ethics Committee (ERN_12-0971P) and performed in accordance with the Declaration of Helsinki.

**Table 1.**
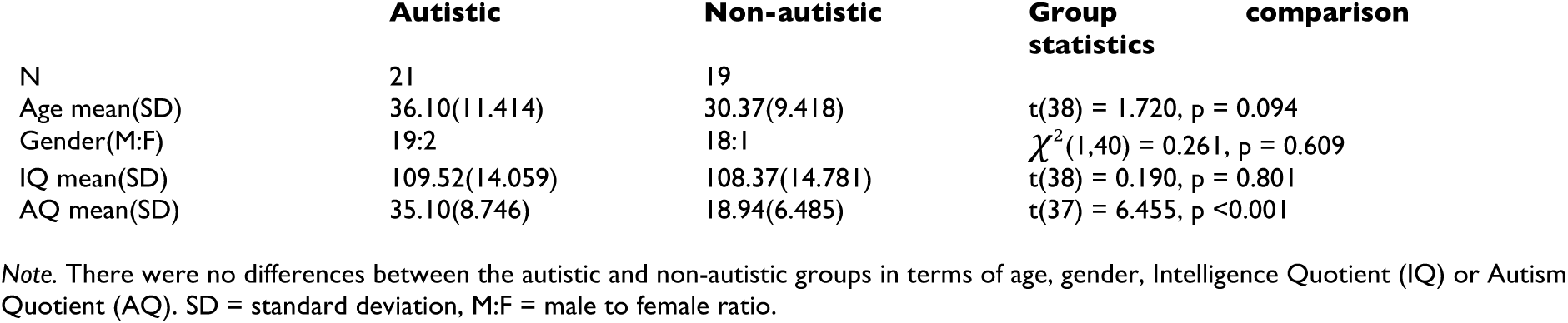
Group Demographics.

### Procedure

A commercial tablet device (Wacom Cintiq 22HD, sampling rate 133Hz, accuracy +/-0.5mm) was used to record movement trajectories. Participants traced shapes displayed on the screen of the tablet with a stylus digitizer in their dominant hand. There were seven conditions relating to seven shapes with different angular frequencies (**Fig 1**). Each trial commenced with written instructions. Participants placed the stylus near the centre of the screen, on a red arrow for spiral condition, or at the darkest part of a blue line for shapes. All movements were anticlockwise and participants were encouraged to draw the shapes fluidly without making corrections. Trials began when participants initiated their drawing movements and finished after they had completed 10 tracings where one tracing was achieved when the number of curvature events equalled the angular frequency (e.g., two curvature events for an ellipse, three for a rounded-triangle), equivalent to a full cycle in shapes with angular frequency >2. Participants undertook at least five trials for each condition. Participants could choose to repeat, or abandon and restart, a trial in which they felt fluidity was not achieved. Hence some subjects undertook more than five trials.

### Data Analysis

All data were processed and analysed in MATLAB R2022a and the data and code required to reproduce the results and figures for this study is freely available at (https://osf.io/j4ncd/?view_only=db95e4126bd14a9bb5aa5401f81d8e7c).

#### Pre-processing

Trials with discontinuous data (pauses or removals of the stylus from the screen) were rejected such that only high-quality trials were included. For each participant the first five high-quality trials were analysed, trials in excess of five were rejected to guard against over-rehearsal effects. This procedure resulted in five trials (where each trial included 10 tracings) for each of the seven conditions.

Error values (i.e., deviation from the ‘correct’ shape) for each trial were calculated as the absolute mean, of the normal distance to the tangent of the nearest point on the shape’s curve as in Madirolas et al.^52^. It should be noted that speed-curvature gradient values (see below for calculation) are highly sensitive to errors (deviations from the ideal shape and the corresponding corrective movement). Thus, if autistic and non-autistic individuals differ in the number of errors this could create an illusory difference in speed-curvature gradients. A linear mixed model predicting error with group (fixed effect) and with trial number and participant ID as random effects demonstrated no significant differences between autistic and non-autistic participants in terms of drawing errors (F(1, 1297) = 2.30, p = .129). Therefore, task compliance is judged to be equivalent between the two groups.

#### Calculation of speed-curvature gradients

To explore the spectrum of power laws, data were analysed according to the procedure outlined in Huh & Sejnowski^25^.

The relationship between tangential speed *ν* and curvature *κ*, as a function of *θ* (in a modified Frenet Serret frame), is given by the following power law (where *φ* = angular frequency of the shape):

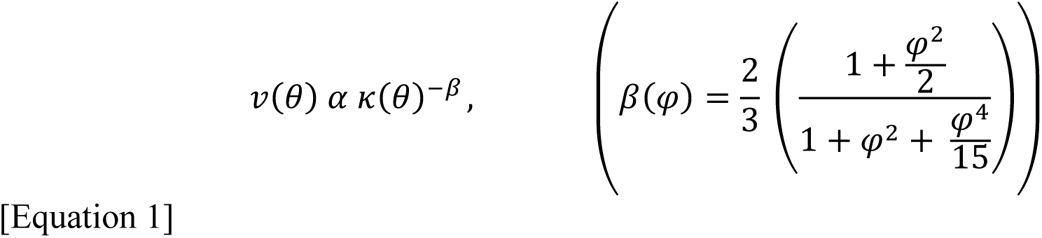

In Equation 1 the value of *β* comprises the gradient of the relationship between (log) speed and (log) curvature. To estimate *β* for each participant, for each condition, the following steps were taken. First the velocities, as a function of *t*, in *x* and *y* directions were obtained by smoothly differentiating the raw positional data.^53^ To express speed and curvature as a function of *θ* state the velocity in the rectangular form *V*(*t*) = *x* + *iy*. Where *x* and *y* are the velocities in the *x* and *y* direction. Which can be re-expressed in the polar form as *V*(*θ*) = *νe*^iθ^ where the radial co-ordinate *ν* = √(*x*^2^ + *y*^2^), i.e., the tangential speed *ν* = |*V* |, and the angular co-ordinate *θ* = tan^−1^(*y*/*x*), the counter clockwise angle from the x-axis. If one restates the right-hand side; *νe*^iθ^ = *e*^iθ+ln(v)^ and takes the natural logarithm to give ln (*V*) = *iθ* +ln (*ν*), one can observe *θ* is the imaginary part of ln(*V*) and tangential speed *ν* is the exponential of the real component of ln(*V*). To ensure monotonicity of *θ*, unwrap *θ*. Smoothly differentiating this *θ* gives angular speed, which divided by previously calculated speed *ν,* gives curvature *κ*. Log-speed *ν* and the log-curvature *κ* profiles were thus obtained and resampled with uniform step-size in *θ*, after spline-fitting. These profiles were low-pass filtered to reject angular frequencies above 25, and subsequently Gaussian band pass filtered around the target angular frequency 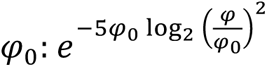. *β* values were obtained by regressing bandpass filtered log-speed against log-curvature.

Our dependent variable was the absolute value of the ‘*β* exponent’. This value corresponds to the gradient of the relationship between log(speed) and log(curvature). A higher *β* thus means a steeper slope of the log curvature vs. log speed relationship (see **Fig. 2** for example for single participant raw data and corresponding *β* estimates). Based on previous findings^25^ we predicted that, for non-autistic participants, there would be a negative relationship between angular frequency of shape (i.e., condition) and *β* value.

**Figure 2.**
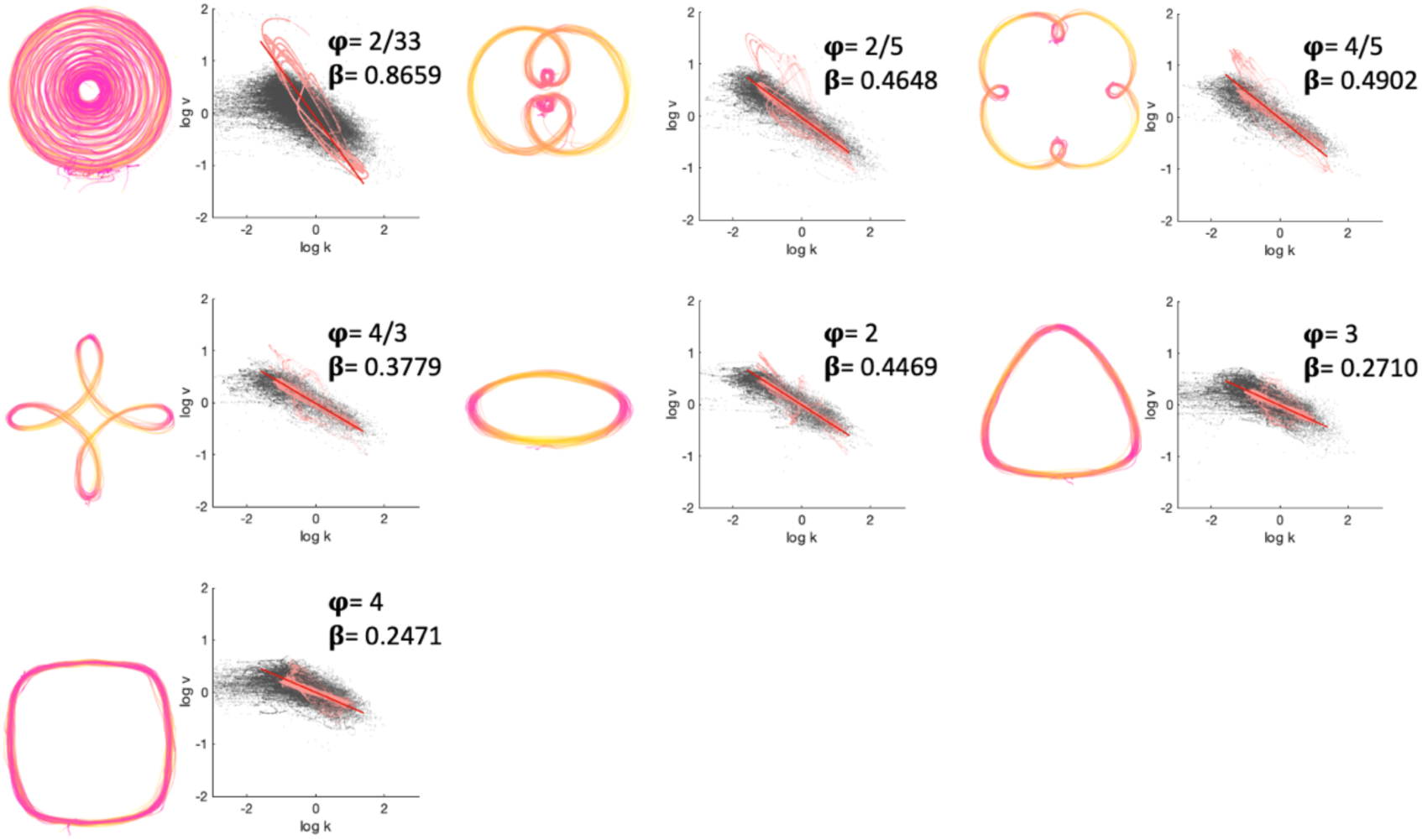
Typical Movement Examples from a Single Participant. The colour scheme represents speed from dark pink (low speed) to yellow (high speed). Log speed vs. log curvature is shown to the right of each shape. Grey dots: Raw curvature-speed data. Pink dots: Bandpass filtered curvature-speed data. Red line: Line of best fit. Measured β exponent is given in the inset for each shape.

An effects-coded, mixed effects model was fitted (using the *fitlme* function of the Statistics and Machine Learning Toolbox in MATLAB) with speed-curvature gradient (absolute value) as the dependent variable (DV) and group (autistic, non-autistic), condition (angular frequency-defined shape 2/33, 2/5, 4/5, 4/3, 2, 3, 4), and the interaction between condition and group as fixed effects. Group and condition were specified as categorical predictors. A random intercept for trial number (1,2,3,4,5) was fitted to account for practice or fatigue effects (e.g., speeding up as a function of trial) and a random intercept for participant (defined as a categorical predictor) was included. To obtain *F* and *p* values we employed analysis of variance for linear mixed effects models using the MATLAB ANOVA function.

#### Frequency analysis

The speed fluctuations that characterise a single drawing movement can be thought of as a complex wave. As with other complex waves - like sound waves - one can use Fourier transform to decompose the wave and represent it as density within different frequency bands. A high-pitched sound wave, for example, would have high spectral density in high frequency bands. Similarly, a drawing of a high angular frequency shape (such as a square) would have a density peak in the band around an angular frequency of four and low density in other bands. Thus, Fourier transform of the speed wave allows one to quantify the relative preponderance of different angular frequencies that is characteristic of a particular drawing.

To plot amplitude spectral density as a function of angular frequency, log speed was examined. The peaks’ periodicity, in every 2π radians, allow the determination of angular frequencies present in the movements of the stylus. Asymmetrical fast Fourier transform (FFT) was employed, which returned the amplitude spectral density of all angular frequencies. Angular frequency values returned by the FFT were then interpolated to obtain uniformly sampled values at 1001 arbitrary points.

Since the FFT assumes an infinite signal, when addressing a finite sample such as the log speed here, the first and last values of each sample must be continuous to avoid artefacts in the FFT results. We addressed this and any general drift in the signal (e.g., from participants generally slowing their movements due to fatigue) by removing a second order polynomial trend.

#### Exploratory analyses of speed, jerk and submovements

For further exploratory analyses (see Supp. Mats. 2) we calculated jerk, speed and number of submovements. In replication of previous analyses,^14,16^ speed for each trial was calculated as the square root of the sum of the squared delta of the x and y vectors. Vectors were low pass filtered at 5 Hz. Jerk was calculated as the second order differential of these vectors. Absolute speed and jerk were averaged, across all samples within a trial for each trial, for each condition, for each participant. Our submovements DV was calculated as the percentage of frames in the whole movement that comprise an acceleration sign change (i.e., a flip from accelerating to decelerating).

## Results

### Autistic movements differ across the spectrum of power law speed-curvature gradients

For speed-curvature gradients, an ANOVA conducted on model coefficients from our mixed model (speed-curvature gradient ~ Condition + Group + (Condition x Group) + (1|Trial) + (1|Participant)); revealed that there was a main effect of group (*F*(1,1285) = 5.29, *p* = .021), main effect of condition (*F*(6,1285) = 1206.40, *p* < .001) and group x condition interaction (*F*(6,1285) = 2.62, *p* = .016; **Fig. 3**). As can be seen from **Table 2** the grand mean is 0.44 (intercept), the estimated mean for the autistic group is 0.46 (grand mean + autistic beta = 0.44 + 0.02), and the estimated mean for the non-autistic group is 0.42 (grand mean + non-autistic beta = 0.44 - 0.02). The main effect of group thus indicates that autistic participants exhibited steeper speed-curvature gradients than non-autistic participants. The main effect of condition indicates that speed-curvature gradients decreased as a function of angular frequency; compared to the grand mean, low angular frequency shapes were characterised by steeper gradients (positive *β* estimates in **Table 2**) and higher angular frequency shapes were characterised by shallower gradients (negative *β* estimates in **Table 2**).

**Figure 3.**
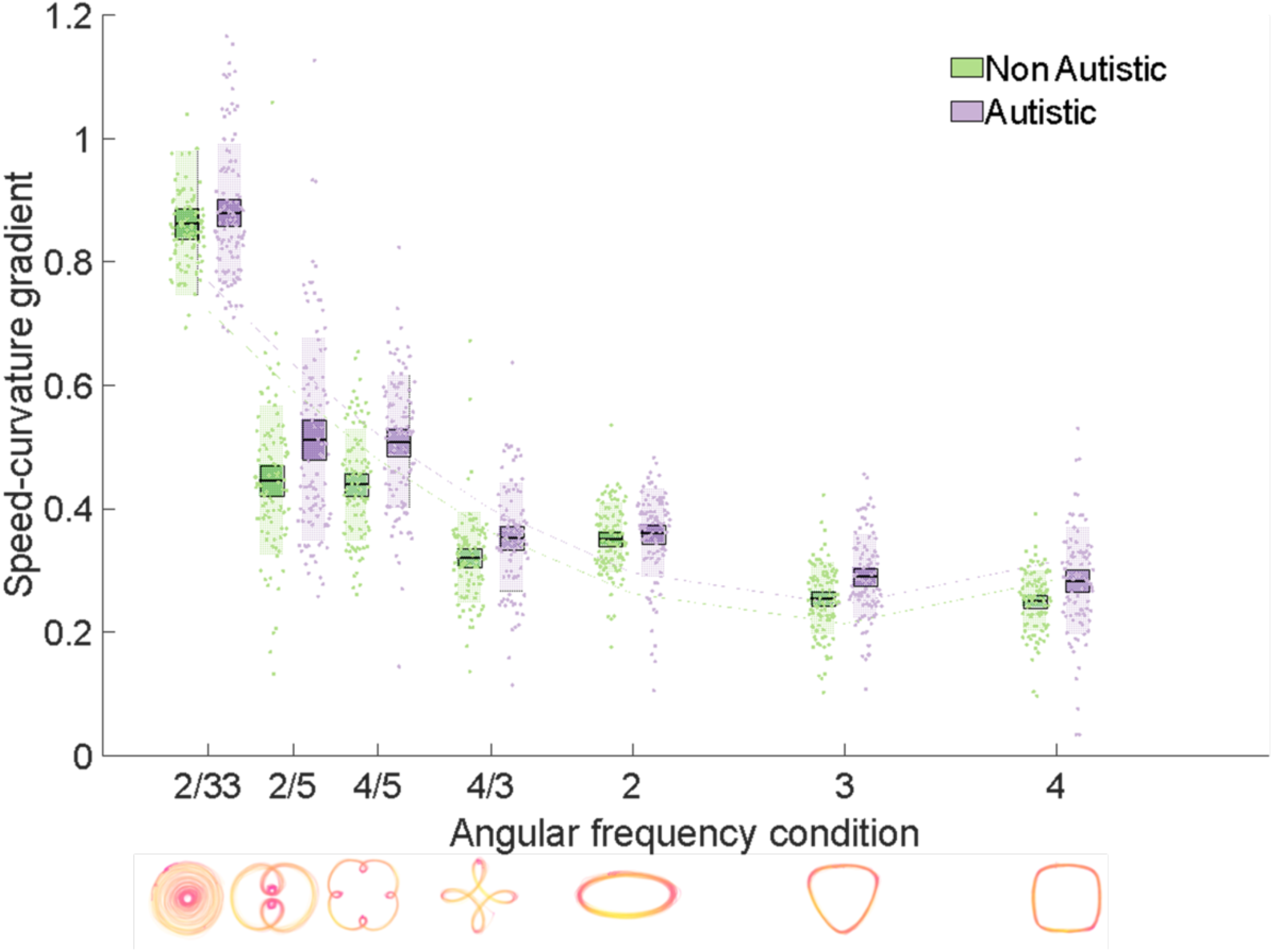
A Graph Depicting Speed-Curvature Gradients for Autistic and Non-autistic Groups. Speed-curvature gradients plotted against angular frequency, for autistic and non-autistic groups. Speed-curvature gradients were significantly negatively related to angular frequency for both autistic and non-autistic groups. However, a main effect of group indicated that speed-curvature gradients were significantly higher for the autistic relative to the non-autistic group, across all levels of angular frequency. Bars = mean, box = standard error of the mean (SEM), individual data points plotted, second order polynomial line of best fit plotted for illustration purposes.

**Table 2.** Model Parameters for Speed-Curvature Gradient Mixed Model.

| | $\beta$ estimate | SE | t statistic | DoF | p value | Lower CI | Upper CI |
| --- | --- | --- | --- | --- | --- | --- | --- |
| **Speed-curvature gradient** |  |  |  |  |  |  |  |
| Intercept | 0.44 | 0.01 | 51.05 | 1285 | <.001*** | 0.42 | 0.45 |
| Non-autistic | -0.02 | 0.01 | -2.30 | 1285 | .022* | -0.04 | 0.00 |
| Autistic | 0.02 | 0.01 | 2.30 | 1285 | .022* | 0.00 | 0.04 |
| 2/33 | 0.43 | 0.01 | 78.65 | 1285 | <.001*** | 0.42 | 0.44 |
| 2/5 | 0.04 | 0.01 | 7.30 | 1285 | <.001*** | 0.03 | 0.05 |
| 4/5 | 0.04 | 0.01 | 6.28 | 1285 | <.001*** | 0.02 | 0.05 |
| 4/3 | -0.10 | 0.01 | -17.64 | 1285 | <.001*** | -0.11 | -0.09 |
| 2 | -0.08 | 0.01 | -14.02 | 1285 | <.001*** | -0.09 | -0.07 |
| 3 | -0.16 | 0.01 | -29.03 | 1285 | <.001*** | -0.17 | -0.15 |
| 4 | -0.17 | 0.01 | -29.42 | 1285 | <.001*** | -0.18 | -0.16 |
| Non-autistic 2/33 | 0.01 | 0.01 | 1.95 | 1285 | .052 | 0.00 | 0.02 |
| Non-autistic 2/5 | -0.01 | 0.01 | -2.32 | 1285 | .020* | -0.02 | 0.00 |
| Non-autistic 4/5 | -0.01 | 0.01 | -2.00 | 1285 | .046* | -0.02 | 0.00 |
| Non-autistic 4/3 | 0.01 | 0.01 | 0.34 | 1285 | .737 | -0.01 | 0.02 |
| Non-autistic 2 | 0.01 | 0.01 | 2.22 | 1285 | .027* | 0.00 | 0.02 |
| Non-autistic 3 | 0.00 | 0.01 | -0.24 | 1285 | .809 | -0.01 | 0.01 |
| Non-autistic 4 | 0.00 | 0.01 | 0.05 | 1285 | .967 | -0.01 | 0.01 |

Note. Model equation: Speed-curvature gradient ~ Condition + Group + (Condition x Group) + (1|Trial) + (1|Participant). Statistics as produced by the MATLAB fitlme function. \*p < .05, \*\*p < .01, \*\*\*p < .001. SE = standard error, DoF = degrees of freedom, CI = confidence interval.

The interaction between group and condition denotes the extra change in the estimated mean gradient over and above the main effects. **Table 2**, for instance, indicates that non-autistic participants for the shape with angular frequency 2/5 have an estimated mean gradient of 0.45 (i.e., the sum of the following *β* coefficients: intercept + non-autistic + ang. freq. 2/5 + non-autistic 2/5 = 0.44 - 0.02 + 0.04 - 0.01 = 0.45). In contrast, the estimated mean gradient for the autistic group for the 2/5 shape is 0.51 (intercept + autistic + ang. freq. 2/5 + autistic 2/5 = 0.44 + 0.02 + 0.04 + 0.01 = 0.51; a difference between the groups of 0.06). *β* coefficients relating to the interaction between group and condition illustrate that the mean estimated difference between groups in speed-curvature gradients varies as a function of condition. Notably, this interaction term is statistically significant for conditions 2/5, 4/5 and 2. For conditions 2/5 and 4/5, the difference between the groups (i.e., higher gradient in the autistic than non-autistic group) is greater than the difference that would be predicted from the main effects of group and condition alone (see **Fig. 3**). For angular frequency 2, the difference between the groups is smaller than the difference that would be predicted from the main effects alone.

A control analysis demonstrated that there was no significant main effect of group (*F*(1,1285) = 0.17, *p* = .684) or interaction between group and condition (*F*(6,1285) = 1.92, *p* = .074) with respect to the fit of the speed-curvature regression line to the data (indexed in terms of log transformed mean squared error values). Thus, the difference between the groups in speed-curvature gradients represents a difference in the gradient of the relationship between log(*ν*) and log(*κ*), and not a difference in the fit of the regression line. Further exploratory mixed models (Supp. Mats. 1) revealed that the groups differed in terms of minimum and maximum speed values, but not curvature values. Autistic participants, compared to non-autistic participants, exhibited lower minimum speeds across all angular frequencies and generally reached higher maximum speeds (though note the difference between the groups in maximum speed was greatest at low angular frequencies). These post hoc analyses demonstrate that differences in speed-curvature gradients between autistic and non-autistic groups are not due to drawing more or less curved movement trajectories (i.e., they are not spatial differences) but rather are due to reaching higher maximum speeds, and slowing to lower minimum speeds.

Further exploratory mixed models (see Supp. Mats. 2 for details) compared the groups in terms of jerk (change in acceleration), speed and number of submovements. Compared to the non-autistic group, autistic participants executed more jerky movements at higher angular frequencies (condition x group interaction: *F*(6, 1285) = 18.24, *p* < .001). At high, but not low, angular frequencies, autistic participants moved more quickly than non-autistic participants (condition x group interaction: *F*(6,1285) = 18.13, *p* < .001). There was also a significant group x condition interaction for submovements (*F*(6, 1285) = 6.31, *p* < .001) indicating that autistic participants executed a greater number of submovements particularly for elliptical shapes. Note that a linear mixed model, with jerk (log transformed) as DV and speed (log transformed), percentage submovements and speed-curvature gradient (all z-scored) as predictors within the same model (with trial and participant included as random factors and condition and group as fixed factors), revealed that speed (*F*(1,1288) = 2036.70, *p* < .001), submovements (*F*(1,1288) = 277.30, *p* < .001) and speed-curvature gradient (*F*(1,1288) = 10.03, *p* = .002) all significantly explained unique variance in jerk values. Thus, indicating that atypically jerky movements in autism are likely due to multiple factors including differences in the way that speed is modulated as a function of curvature as well as overall differences in mean speed, and the number of submovements that individuals execute.

### FFT of speed offers insight into mechanisms underpinning high speed-curvature gradients in autism

A drawing of an ellipse should have a clearly defined density peak in a band centred around angular frequency 2 (because there are two notable changes in curvature in an ellipse) and low density in other bands (see **Fig. 4** left panel for example amplitude spectral density function for the angular frequency 2 shape). We employed FFT to provide further insight into the atypically high speed-curvature gradients in the autistic group. Specifically, we wondered whether autistic participants were modulating their speed in a way that is appropriate for drawing a different shape. For instance, if participants modulated speed in a way that is appropriate for drawing a triangle when actually drawing a square this would result in an abnormally high speed-curvature gradient (because speed-curvature gradient increases with decreasing angular frequency) and would be revealed by a leftward-shifted peak in the FFT-derived amplitude spectral density function. To investigate this possibility, we calculated amplitude spectral density functions for all participants, for all angular frequency-defined shapes. Visual exploration of the data clearly showed that amplitude spectra were not leftward-shifted for the autistic group; in fact, peaks were almost exactly aligned. Nevertheless, we observed that peaks were more precise (more narrow) for non-autistic relative to autistic participants. To quantify this, we aligned amplitude spectra along the target frequency (**Fig. 4** right panel) such that the zero point corresponds to an angular frequency of two for drawings of elliptical shapes, three for the triangle shape, four for the square shape and so on. To statistically compare the profiles, we used t-tests at all 1001 points on the angular frequency spectrum.

**Figure 4.**
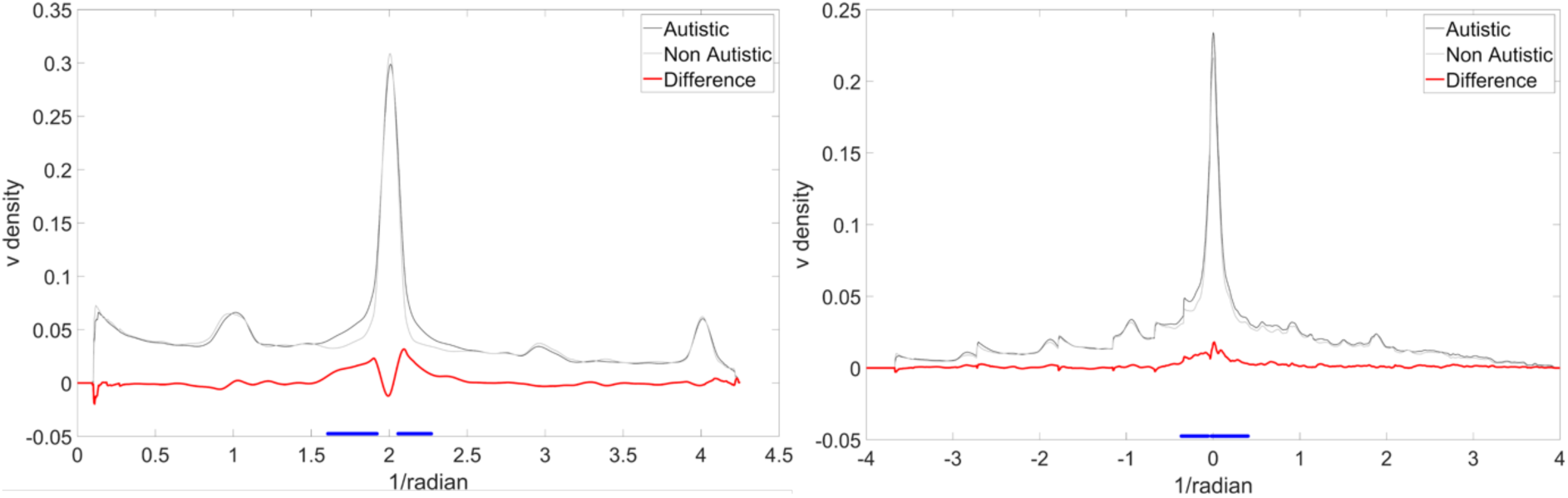
Amplitude Spectral Density Graphs. Left panel: Illustrative density spectrum for angular frequency 2. Note the clear peak at 2, which is more precise (narrower peak flanks) for the non-autistic group. Right panel: Amplitude spectral density of velocity for all angular frequency-defined conditions aligned to zero. Black line = autistic, Grey line = non-autistic, Red line = autistic - non-autistic, blue lines denote bootstrapped clusters of statistical significance.

This analysis revealed that autistic participants exhibited reduced precision of speed oscillations around the target frequency. Bootstrapped *t*-tests (1000 boots) showed two significant clusters of difference - defined as clusters of difference that occurred in less than 5% of comparisons with resampled distributions (highlighted in blue in **Fig. 4** right panel) - between the autistic and non-autistic groups. These clusters fell either side of the target frequency, thus statistically confirming the reduced precision of speed oscillations around the target frequency for the autistic group. It should be noted that this analysis was repeated for curvature and no significant differences were observed, corroborating matched task compliance between the two groups (Supp. Mats. 3).

Since our bootstrapped analysis does not account for repeated observations from the same participants across multiple trials we used trapezoidal numerical integration (MATLAB trapz.m) to calculate the area under the amplitude spectral density function (FFT integral) for all trials, for all participants. FFT integrals were submitted to an effects-coded, mixed effects model with condition, group and the interaction between condition and group as fixed effects and trial and participant as random effects (FFT integral ~ Condition + Group + (Condition x Group) + (1|Trial) + (1|Participant); **Table 3**). An ANOVA conducted on model coefficients revealed that there was a significant main effect of group (*F*(1,1283) = 4.63, *p* = .032), main effect of condition (*F*(6,1283) = 357.19, *p* < .001) and group x condition interaction (*F*(6,1283) = 2.52, *p* = .020). The main effect of group indicates that the FFT integral was greater for autistic relative to non-autistic participants and the interaction demonstrates that this difference varied as a function of angular frequency, with the difference being small for the 2/33 condition and particularly large at an angular frequency of 4. In sum, we observed that non-autistic participants showed precise (narrow and tall) peaks around target frequencies whereas density spectra for autistic participants were less precise (wider and shorter) (**Fig. 5**).

**Figure 5.**
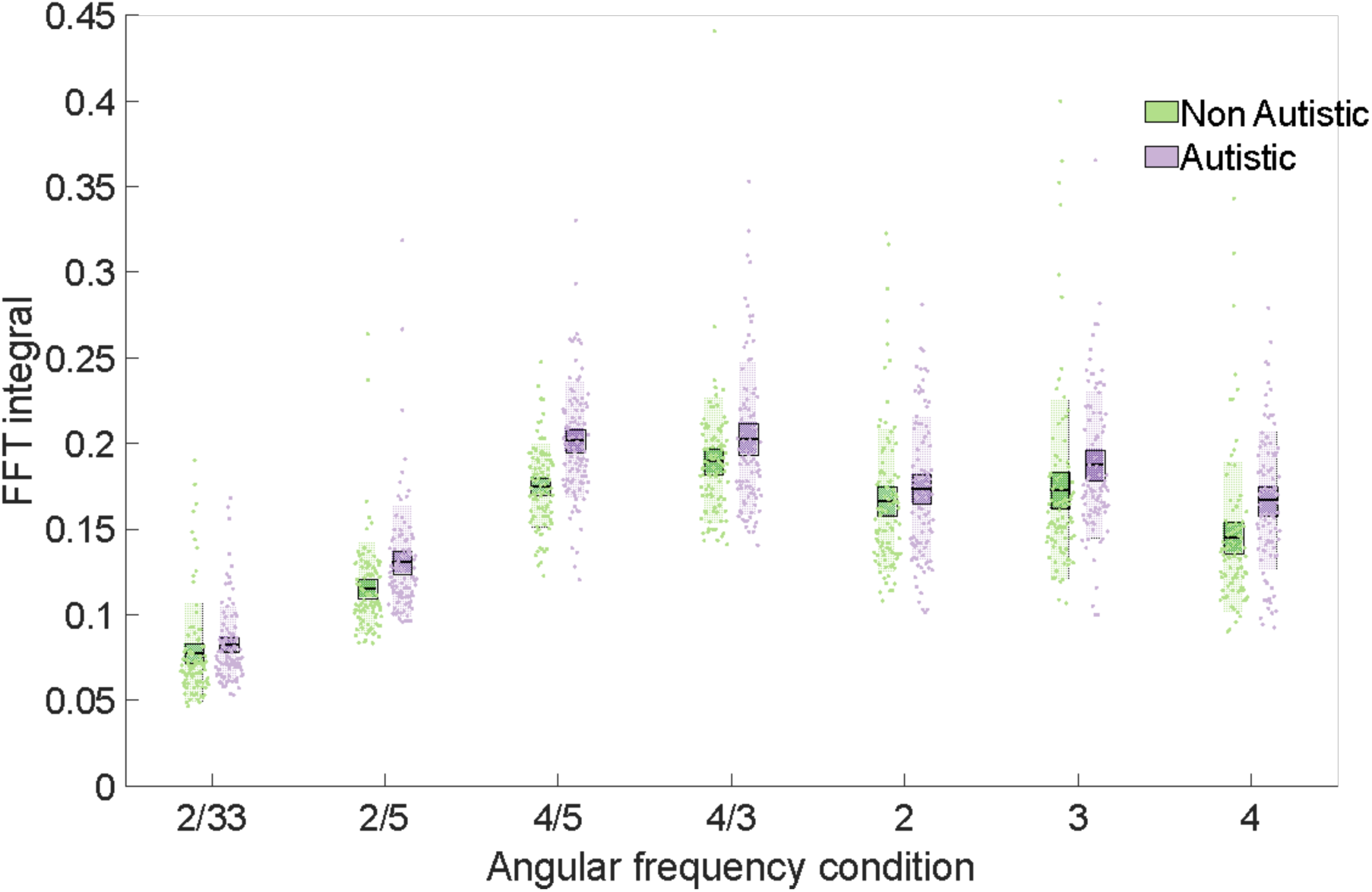
A Graph Depicting the FFT integrals for Autistic and Non-autistic Groups. FFT integral against angular frequency for autistic and non-autistic groups. FFT integral values were significantly higher for the autistic relative to the non-autistic group and this difference was particularly large for the 4/5 condition. Bars = mean, box = SEM, individual data points plotted.

**Table 3.** Model Parameters for FFT Integral Mixed Model.

| | $\beta$ estimate | SE | t statistic | DoF | p value | Lower CI | Upper CI |
| --- | --- | --- | --- | --- | --- | --- | --- |
| **FFT integral** |  |  |  |  |  |  |  |
| Intercept | 0.16 | 0.00 | 42.40 | 1283 | <.001*** | 0.15 | 0.16 |
| Non-autistic | -0.01 | 0.00 | -2.15 | 1283 | .032* | -0.02 | 0.00 |
| Autistic | 0.01 | 0.00 | 2.15 | 1283 | .032* | 0.00 | 0.02 |
| 2/33 | -0.08 | 0.00 | -37.69 | 1283 | <.001*** | -0.08 | -0.07 |
| 2/5 | -0.03 | 0.00 | -16.77 | 1283 | <.001*** | -0.04 | -0.03 |
| 4/5 | 0.03 | 0.00 | 16.30 | 1283 | <.001*** | 0.03 | 0.04 |
| 4/3 | 0.04 | 0.00 | 18.96 | 1283 | <.001*** | 0.04 | 0.04 |
| 2 | 0.01 | 0.00 | 6.68 | 1283 | <.001*** | 0.01 | 0.02 |
| 3 | 0.02 | 0.00 | 11.52 | 1283 | <.001*** | 0.02 | 0.03 |
| 4 | 0.00 | 0.00 | -0.12 | 1283 | .908 | 0.00 | 0.00 |
| Non-autistic 2/33 | 0.01 | 0.00 | 2.91 | 1283 | .004** | 0.00 | 0.01 |
| Non-autistic 2/5 | 0.00 | 0.00 | -0.11 | 1283 | .912 | 0.00 | 0.00 |
| Non-autistic 4/5 | 0.00 | 0.00 | -1.66 | 1283 | .097 | -0.01 | 0.00 |
| Non-autistic 4/3 | 0.00 | 0.00 | 0.05 | 1283 | .959 | 0.00 | 0.00 |
| Non-autistic 2 | 0.00 | 0.00 | 1.41 | 1283 | .160 | 0.00 | 0.01 |
| Non-autistic 3 | 0.00 | 0.00 | -0.50 | 1283 | .620 | -0.01 | 0.00 |
| Non-autistic 4 | 0.00 | 0.00 | -2.03 | 1283 | .043* | -0.01 | 0.00 |

Note. Model equation: FFT integral ~ Condition + Group + (Condition x Group) + (1|Trial) + (1|Participant). Statistics as produced by the MATLAB fitlme function. \* $p < .05$ , \*\* $p < .01$ , \*\*\* $p < .001$ . SE = standard error, DoF = degrees of freedom, CI = confidence interval.

## Discussion

The current study investigated whether movements executed by autistic adults diverge from the typical power law relationship linking movement speed and curvature. Compared to age-, IQ- and sex-matched non-autistic participants, the gradient of the relationship between speed and curvature was significantly higher for autistic individuals across angular frequency-defined shapes from 2/33 (spiral-like shapes) to 4 (square-like shapes), though the difference between the groups was particularly large for the shapes defined by the angular frequencies 2/5 and 4/5. Post hoc analyses demonstrated that differences in speed-curvature gradients between the groups were not due to spatial differences; autistic participants did not draw more/less curved trajectories. Instead, gradient differences were due to differences in speed modulation. That is, autistic participants’ movement was characterised by accelerating to higher maximum speeds (on the straights) and slowing to lower minimum speeds (at the corners).

Higher speed-curvature gradients in autism may indicate underlying differences in motor control policies and/or biomechanical constraints. The negative relationship between speed and curvature has received much attention for at least two reasons; first, because it is robust across effectors and species, but also because discussion about its origins has shaped theories of the computational principles, and biomechanical constraints, that govern movements. The one third power law aligns remarkably well with control policies that optimise movement smoothness (e.g., minimum-jerk model^17,18^). In other words, if one could *program* a robot to move as smoothly as possible, one would observe that the robot’s movements also exhibit a negative relationship between speed and curvature. It has been theorised that the minimisation of jerk may therefore be a higher-order goal of the human motor system.^17,43,54^ According to this view, departures from the one third power law indicate brain-based differences in motor cortical control policies. Alternative accounts argue that the one third power law is a product of the biomechanics of the body.^34,35^ That is, muscles and joints effectively comprise a system of filters, and even a uniform velocity signal fed through the same filters would be modulated as a function of trajectory curvature. According to this view, departures from the one third power law may indicate differences in biomechanical constraints. Consequently, our data help to direct future mechanistic work concerning motor control in autism. Specifically, higher speed-curvature gradients in autistic participants suggest that future studies need to a) test whether autistic bodies are subject to different biomechanical constraints (which may, for instance, promote movements around certain joints with corresponding power law differences^55^) and b) assess whether autistic movements may be governed by different control policies (e.g., minimising a cost function other than smoothness).

We explored potential reasons that autistic participants might exhibit steeper gradients between speed and curvature by using FFT to decompose the speed profile and represent it as the relative preponderance of speed changes in different angular frequency bands. In contrast to our expectations, we did not find that the autistic participants were modulating speed in a way that is appropriate for drawing a different shape (e.g., modulating speed in a way that is appropriate for drawing a triangle when drawing a square). Instead, we observed that whilst non-autistic participants showed precise (narrow) peaks around target frequencies, density spectra for autistic participants were less precise (broader). Interpretation of our results may be informed by auditory science. The FFT operation that we here carried out on the speed wave is analogous to the operation that is carried out on sound waves by the basilar membrane in the human ear. The basilar membrane behaves as if it contains a bank of overlapping bandpass filters (‘auditory filters’), which decompose sound waves into their constituent frequency components. Each filter is tuned to a particular centre frequency, with that basilar membrane filter responding maximally to that frequency and progressively less to more distal frequencies. The relative response of the filter, as a function of frequency, is known as the auditory filter shape and is analogous to the amplitude spectral density function that we calculated for each angular frequency-defined shape. In normal hearing participants auditory filters are sharply tuned around the centre frequency.^56^ However, Plaisted and colleagues^57^ reported atypically broad auditory filters in autistic individuals, wherein the bandwidth of the filter was significantly wider than bandwidths measured in non-autistic individuals.^58^ Plaisted and colleagues^57^ suggest that their results indicate differences in the peripheral filtering of incoming sensory signals, as opposed to indicating differences in central mechanisms (though they note that the two are not mutually exclusive). Plaisted and colleagues’^57^ interpretation is highly relevant for our results. The appropriate modulation of speed as a function of curvature has been argued to, at least in part, comprise the end product of a series of filters,^34^ which can tune a noisy motor signal to the appropriate target (angular) frequency. Thus, mirroring Plaisted and colleagues’^57^ results, which may indicate differences in the filtering of *incoming* sensory signals, our results may indicate differences in the filtering of *outgoing* motor signals. Clearly, further studies are required to empirically test this hypothesis, and arbitrate between models that postulate atypical filtering of motor signals in autism and models that postulate that the initial motor signal is atypically noisy.^22^

The current results have practical implications for the early identification of autism. Tools that can discriminate between autistic and non-autistic individuals from an early age are much sought after^59^ and movement-based methods are particularly appealing because they do not rely on language and can therefore be used with non-verbal children.^60^ Identification of autism can also be difficult where camouflaging is common, for example in females^61^ and those with higher cognitive and executive function abilities.^62,63^ Movement atypicalities may be less susceptible to camouflaging, perhaps owing to differences between individuals being more subtle, and explicit rules about typical motion being harder to identify, meaning movement-based methods might also prove effective for discriminating between autistic and non-autistic adults who may otherwise perform typically on measures assessing social and communication difficulties or restricted interests and repetitive behaviours. Here we showed that differences between autistic and non-autistic groups are particularly large for the shapes defined by the angular frequencies 2/5 and 4/5, thus suggesting that the speed-curvature gradient relating to these particular trajectories may be usefully incorporated into automated tools.^26^ Since speed-curvature gradients are mean speed- and scale-invariant^25^ they have the added advantage that they are robust against differences in speed and scale that are known to affect autistic movement but which do not systematically discriminate between autistic and non-autistic individuals (i.e., handwriting analysis has linked autism to both micro- and macro-graphia).^8,9,64^ Further studies, however, are required to investigate whether our results can be replicated in young autistic children, and to test whether speed-curvature gradients can discriminate between autism and other developmental conditions such as developmental coordination disorder.

Our results may also feed into the development of motor control support systems for autistic individuals. By asking our participants to draw shapes which span the angular frequency spectrum, we have been able to show that autistic individuals exhibit atypical speed-curvature gradients across the spectrum. In principle, the shapes we asked participants to draw can be combined to create an infinite number of trajectory shapes. Conversely any trajectory (even a seemingly random doodle) can be decomposed into density across the angular frequency spectrum.^25,30^ Our results therefore suggest that autistic individuals will likely exhibit atypical speed-curvature gradients that extend beyond drawing simple shapes and perhaps even encompass functional tasks such as handwriting or throwing. Indeed, previous studies have shown that minimally-jerky movement, that accords with the fundamental power laws, is important across a range of functional motor tasks including handwriting^65,66^ and throwing^17,18,67^, and that autistic individuals often experience difficulties with such tasks.^7–11^ It is possible that the atypical speed-curvature relationship that we have observed underpins these functional challenges. Ongoing endeavours to train movement kinematics^68^ have the potential to reveal novel methods which may be used to support autistic people in performing functional motor tasks.

In addition to speed-curvature differences, we replicated previously reported differences in movement jerk in autism^16,21,22^ and observed that, for higher angular frequency shapes, autistic participants decomposed shapes into a greater number of submovements and moved faster. Further studies are needed to provide mechanistic explanations for atypical movement speed in autism. Such studies may, for instance, test whether autistic movements are governed by an atypical subjective cost function, as has been suggested for the atypically slow movements seen in people with Parkinson’s disease.^40^ It is similarly unknown why autistic individuals would decompose movements into a greater number of submovements. An interesting hypothesis here - with roots in cognitive theories of autism including weak central coherence^69^ and enhanced perceptual functioning^70^ that propose a preference towards local over global processing - is that autistic individuals may decompose actions into a piecemeal series of smaller individual steps.^4,44,71–74^ Importantly, our linear mixed model indicated that speed-curvature gradients, speed and submovements all predicted unique variance in jerk. Thus, atypically jerky movements in autism are likely due to multiple factors including deviations from the typical power law function as well as overall differences in mean speed, and the number of submovements that individuals execute. Consequently, there may be multiple underlying mechanisms that, when combined together, explain the jerky profile of movements that is commonly observed in autism.

To conclude, our results evidence, in autistic adults, a deviation from the power laws that typically govern movement and suggest atypical filtering of outgoing movement signals as evidenced by the reduced precision of speed oscillations around the target frequency. These data raise important questions about motor control policies and biomechanical constraints that govern autistic movement and may have important implications for identification and lifelong support systems for autistic people.

## Data availability statement

Data and analysis are available online at the following repository https://osf.io/j4ncd/?view_only=db95e4126bd14a9bb5aa5401f81d8e7c

## Acknowledgements

The authors would like to thank Bianca Schuster for data analysis advice and the Birmingham Participatory Autism Research Team (B-PART) consultancy group for providing advice, from the perspective of individuals with lived experience of autism, regarding the reporting of these findings.

## Supplementary Material 1

### Analyses of minimum and maximum speed

Exploratory analyses were conducted on the minimum and maximum speed and curvature values, using the mixed model formula: DV ~ Condition + Group + (Condition x Group) + (1|Trial) + (1|Participant). The following transforms were used on the data: log (max velocity), reciprocal (max curvature (see below)), reflect reciprocal (min curvature, min velocity). Analyses revealed that the groups differed in terms of minimum and maximum speed values. For maximum speed there was a significant main effect of condition (*F*(6,1285) = 61.85, *p* < .001) and an interaction between group and condition (*F*(6,1285) = 2.68, *p* = .014), indicating that autistic participants generally reached higher maximum speeds but that the difference between the groups was greatest at low, compared to higher angular frequencies. There was no significant main effect of group (*F*(1,1285) = 3.08, *p* = .079). For minimum speed there was a significant main effect of group (*F*(1,1285) = 8.31, *p* = .004), and condition (*F*(6, 1285) = 4.78, *p* < .001), and no interaction between group and condition (*F*(6, 1285) = 0.91, *p* = .483). Examination of *β* estimates in **Supplementary Table 1** indicates that autistic participants generally exhibited reduced minimum speeds (i.e., a negative *β* estimate) compared to non-autistic participants.

In contrast, the groups did not differ in terms of their minimum and maximum curvature values. For maximum curvature, the main effect of group was not significant (*F*(1, 1285) = 1.74, *p* = .187), nor was the interaction between group and condition (*F*(6, 1285) = 0.90, *p* = .493). For minimum curvature, non-significant results were found for both the main effect of group (*F*(1, 1285) = 1.52, *p* = .218) and the interaction between group and condition (*F*(6, 1285) = 1.58, *p* = .150).

**Supplementary Table 1.** Model Parameters for Maximum and Minimum Speed Mixed Models.

| | $\beta$ estimate | SE | t statistic | DoF | p value | Lower CI | Upper CI |
| --- | --- | --- | --- | --- | --- | --- | --- |
| **Maximum Speed** |  |  |  |  |  |  |  |
| Intercept | -0.30 | 0.02 | -18.90 | 1285 | <.001*** | -0.33 | -0.27 |
| Non-autistic | -0.03 | 0.01 | -1.76 | 1285 | .079 | -0.05 | 0.00 |
| Autistic | 0.03 | 0.01 | 1.76 | 1285 | .079 | 0.00 | 0.05 |
| 2/33 | 0.04 | 0.01 | 3.23 | 1285 | .001** | 0.01 | 0.06 |
| 2/5 | 0.15 | 0.01 | 12.71 | 1285 | <.001*** | 0.12 | 0.17 |
| 4/5 | 0.04 | 0.01 | 3.69 | 1285 | <.001*** | 0.02 | 0.07 |
| 4/3 | -0.02 | 0.01 | -1.32 | 1285 | .186 | -0.04 | 0.01 |
| 2 | 0.02 | 0.01 | 1.86 | 1285 | .063 | 0.00 | 0.05 |
| 3 | -0.06 | 0.01 | -4.67 | 1285 | <.001*** | -0.08 | -0.03 |
| 4 | -0.18 | 0.01 | -14.99 | 1285 | <.001*** | -0.21 | -0.16 |
| Non-autistic 2/33 | -0.01 | 0.01 | -1.27 | 1285 | .204 | -0.04 | 0.01 |
| Non-autistic 2/5 | -0.01 | 0.01 | -1.01 | 1285 | .311 | -0.03 | 0.01 |
| Non-autistic 4/5 | -0.02 | 0.01 | -1.34 | 1285 | .181 | -0.04 | 0.01 |
| Non-autistic 4/3 | 0.03 | 0.01 | 2.29 | 1285 | .022* | 0.01 | 0.05 |
| Non-autistic 2 | 0.03 | 0.01 | 2.65 | 1285 | .008** | 0.01 | 0.06 |
| Non-autistic 3 | 0.00 | 0.01 | 0.13 | 1285 | .899 | -0.02 | 0.03 |
| Non-autistic 4 | -0.01 | 0.01 | -1.48 | 1285 | .140 | -0.04 | 0.01 |
| **Minimum Speed** |  |  |  |  |  |  |  |
| Intercept | 0.59 | 0.01 | 74.31 | 1285 | <.001*** | 0.58 | 0.61 |
| Non-autistic | 0.02 | 0.01 | 2.88 | 1285 | .004** | 0.01 | 0.04 |
| Autistic | -0.02 | 0.01 | -2.88 | 1285 | .004** | -0.04 | -0.01 |
| 2/33 | -0.03 | 0.01 | -2.59 | 1285 | .010** | -0.05 | -0.01 |
| 2/5 | -0.04 | 0.01 | -3.80 | 1285 | <.001*** | -0.06 | -0.02 |
| 4/5 | 0.00 | 0.01 | 0.15 | 1285 | .883 | -0.02 | 0.02 |
| 4/3 | 0.00 | 0.01 | 0.44 | 1285 | .661 | -0.02 | 0.03 |
| 2 | 0.03 | 0.01 | 2.71 | 1285 | .007** | 0.01 | 0.05 |
| 3 | 0.01 | 0.01 | 0.89 | 1285 | .372 | -0.01 | 0.03 |
| 4 | 0.02 | 0.01 | 2.05 | 1285 | .041* | 0.00 | 0.04 |
| Non-autistic 2/33 | -0.01 | 0.01 | -1.02 | 1285 | .310 | -0.03 | 0.01 |
| Non-autistic 2/5 | -0.01 | 0.01 | -0.53 | 1285 | .599 | -0.03 | 0.02 |
| Non-autistic 4/5 | 0.01 | 0.01 | 0.47 | 1285 | .639 | -0.02 | 0.03 |
| Non-autistic 4/3 | -0.01 | 0.01 | -1.18 | 1285 | .237 | -0.03 | 0.01 |
| Non-autistic 2 | 0.01 | 0.01 | 1.13 | 1285 | .259 | -0.01 | 0.03 |
| Non-autistic 3 | 0.00 | 0.01 | -0.37 | 1285 | .715 | -0.02 | 0.02 |
| Non-autistic 4 | 0.02 | 0.01 | 1.45 | 1285 | .149 | -0.01 | 0.04 |

Note. Model equation: DV ~ Condition + Group + (Condition x Group) + (1|Trial) + (1|Participant). Statistics as produced by the Matlab fitlme function. \*p < .05, \*\*p < .01, \*\*\*p < .001.

## Supplementary Material 2

### Exploratory analyses: speed, jerk and submovements

Three separate effects coded linear mixed effects models were fitted (using the *fitglme* function of the Statistics and Machine Learning Toolbox in MATLAB) with jerk, submovements and speed as the DVs. Each model included group (autistic, non-autistic), condition (angular frequency 2/33, 2/5, 4/5, 4/3, 2, 3, 4), and the interaction between condition and group as fixed effects. Group and condition were specified as categorical predictors. A random intercept for trial number (1,2,3,4,5) was fitted to account for practice or fatigue effects (e.g., speeding up as a function of trial) and a random intercept for participant (defined as a categorical predictor) was included. The following transforms were used on the data: log (speed, jerk). Where DVs were not normally distributed (i.e., jerk and speed) a logarithmic link function was specified.

#### Autistic hand movements are characterised by increased jerk at higher angular frequencies

An ANOVA conducted on coefficients from the mixed effects model (Jerk ~ Condition + Group + (Condition x Group) + (1|Trial) + (1|Participant),‘Link’, ‘log’) revealed that there was a main effect of condition (*F*(6,1285) = 22.70, *p* < .001) and an interaction between group and condition (*F*(6, 1285) = 18.24, *p* < .001; **Supplementary Fig. 1**). The main effect of group was not significant (*F*(1, 1285) = 3.61, *p* = .058), though there was a numerical difference between groups. As can be seen from **Supplementary Table 2** the grand mean (log(jerk)) is 5.25 (intercept), the estimated mean for the autistic group is 5.45 (grand mean + autistic beta = 5.25 + 0.2), and the estimated mean for the non-autistic group is 5.05 (grand mean + non-autistic beta = 5.25 - 0.2). Thus, relative to the non-autistic group, the autistic group produced numerically more jerky movements. The main effect of condition indicates that jerk increases as a function of angular frequency condition. That is, relative to the grand mean, estimated jerk is lower for conditions 2/33 (*β* = −0.17), 2/5 (*β* = −15), 4/5 (*β* = −0.1) and higher for conditions 4/3 (*β* = 0.07), 2 (*β* = 0.33) and 3 (*β* = 0.08).

The interaction between group and condition denotes the extra change in the estimated mean jerk over and above the main effects. **Supplementary Table 2** indicates that non-autistic participants in the 2/33 condition have an estimated mean jerk of 4.98 (i.e., the sum of the following *β* coefficients: intercept + non-autistic + 2/33 + non-autistic 2/33 = 5.25 - 0.2 - 0.17 + 0.1 = 4.98). In contrast, estimated mean jerk for the autistic group in the 2/33 condition is 5.18 (intercept + autistic + 2/33 + autistic 2/33 = 5.25 + 0.2 - 0.17 - 0.1 = 5.18; a difference between the groups of 0.2). Positive interaction *β*s in **Supplementary Table 2** indicate that the difference between the autistic and non-autistic groups is smaller than that which would be predicted by the main effects of group and condition alone, negative interaction *β*s indicate that the difference is greater than would be predicted by group and condition alone. Thus, the interaction indicates that the difference in jerk between the autistic and non-autistic groups is particularly large at higher angular frequencies (angular frequency defined shapes 2,3 and 4). For example, the biggest difference between the groups is for the 2 angular frequency shape where estimated jerk for the non-autistic group = 5.25 - 0.2 + 0.33 - 0.21 = 5.17, whereas estimated jerk for the autistic group = 5.25 + 0.2 + 0.33 + 0.21 = 5.99 (a difference of 0.82).

**Supplementary Figure 1.**
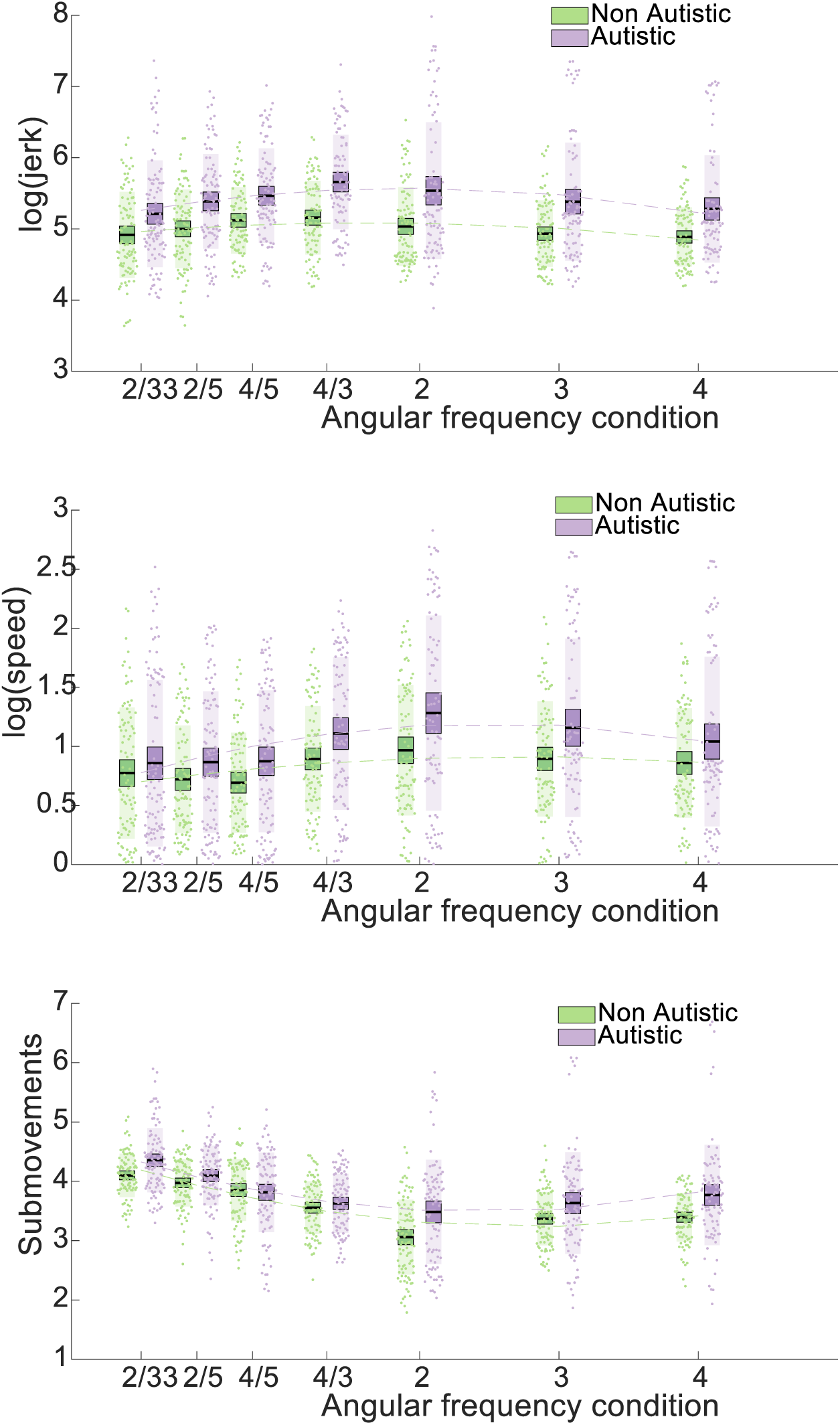
Graphs of Speed, Jerk and Submovements for Autistic and Non-autistic Groups. Log speed and jerk, and submovements plotted against angular frequency (condition) for autistic (purple) and non-autistic (green) groups. Bars = mean, box = SEM, individual data points plotted, second order polynomial line of best fit plotted for illustration purposes.

#### Autistic hand movements are characterised by increased speed at higher angular frequencies

For speed, there was no main effect of Group (*F*(1,1285) = 1.16, *p* = .282). There was, however, a significant main effect of condition (*F*(6,1285) = 107.60, *p* < .001) and group by condition interaction (*F*(6,1285) = 18.13, *p* < .001). Coefficients from a mixed effects model (Speed ~ Condition + Group + (Condition x Group) + (1|Trial) + (1|Participant), ‘Link’, ‘log’) can be seen in **Supplementary Table 2**. The main effect of condition was driven by an increase in speed as a function of angular frequency. Notably, the *β* values (**Supplementary Table 2**) show that, relative to the grand mean, speed is most greatly increased for the angular frequency 2 condition, thus speed and angular frequency are related by an inverted-U-shaped function (**Supplementary Fig. 1**). *β* coefficients relating to the interaction between group and condition illustrate that the mean estimated speed difference between groups varied as a function of condition. Notably, interaction *β*s are positive for low angular frequencies and negative for high angular frequencies. This indicates that, at high angular frequencies the difference in speed between the autistic and non-autistic groups is larger than that which would be predicted from the main effects alone (and smaller than predicted at low angular frequencies).

**Supplementary Table 2.** Model Parameters for Jerk and Speed Mixed Models.

| | $\beta$ estimate | SE | t statistic | DoF | p value | Lower CI | Upper CI |
| --- | --- | --- | --- | --- | --- | --- | --- |
| **Jerk** |  |  |  |  |  |  |  |
| Intercept | 5.25 | 0.11 | 47.82 | 1285 | <.001*** | 5.03 | 5.46 |
| Non-autistic | -0.20 | 0.11 | -1.90 | 1285 | .058 | -0.41 | 0.01 |
| Autistic | 0.20 | 0.11 | 1.90 | 1285 | .058 | -0.01 | 0.41 |
| 2/33 | -0.17 | 0.04 | -4.22 | 1285 | <.001*** | -0.25 | -0.09 |
| 2/5 | -0.15 | 0.04 | -4.08 | 1285 | <.001*** | -0.23 | -0.08 |
| 4/5 | -0.10 | 0.04 | -2.85 | 1285 | .004** | -0.18 | -0.03 |
| 4/3 | 0.07 | 0.03 | 2.23 | 1285 | .025* | 0.01 | 0.14 |
| 2 | 0.33 | 0.03 | 10.24 | 1285 | <.001*** | 0.27 | 0.40 |
| 3 | 0.08 | 0.04 | 2.07 | 1285 | .039* | 0.00 | 0.16 |
| 4 | -0.06 | 0.04 | -1.44 | 1285 | 0.149 | -0.14 | 0.02 |
| Non-autistic 2/33 | 0.10 | 0.04 | 2.60 | 1285 | .009** | 0.03 | 0.18 |
| Non-autistic 2/5 | 0.17 | 0.04 | 4.55 | 1285 | <.001*** | 0.10 | 0.25 |
| Non-autistic 4/5 | 0.17 | 0.04 | 4.51 | 1285 | <.001*** | 0.10 | 0.24 |
| Non-autistic 4/3 | 0.09 | 0.03 | 2.74 | 1285 | .006** | 0.03 | 0.16 |
| Non-autistic 2 | -0.21 | 0.03 | -6.39 | 1285 | <.001*** | -0.27 | -0.14 |
| Non-autistic 3 | -0.18 | 0.04 | -4.72 | 1285 | <.001*** | -0.26 | -0.11 |
| Non-autistic 4 | -0.14 | 0.04 | -3.28 | 1285 | .001** | -0.22 | -0.06 |
| **Speed** |  |  |  |  |  |  |  |
| Intercept | 0.93 | 0.10 | 9.70 | 1285 | <.001*** | 0.74 | 1.12 |
| Non-autistic | -0.10 | 0.10 | -1.08 | 1285 | .282 | -0.29 | 0.09 |
| Autistic | 0.10 | 0.10 | 1.08 | 1285 | .282 | -0.09 | 0.29 |
| 2/33 | -0.09 | 0.02 | -4.59 | 1285 | <.001*** | -0.13 | -0.05 |
| 2/5 | -0.23 | 0.02 | -10.38 | 1285 | <.001*** | -0.27 | -0.18 |
| 4/5 | -0.27 | 0.02 | -11.47 | 1285 | <.001*** | -0.31 | -0.22 |
| 4/3 | 0.00 | 0.02 | 0.02 | 1285 | 0.981 | -0.04 | 0.04 |
| 2 | 0.32 | 0.02 | 20.29 | 1285 | <.001*** | 0.29 | 0.35 |
| 3 | 0.18 | 0.02 | 10.28 | 1285 | <.001*** | 0.14 | 0.21 |
| 4 | 0.09 | 0.02 | 4.61 | 1285 | <.001*** | 0.05 | 0.12 |
| Non-autistic 2/33 | 0.06 | 0.02 | 2.97 | 1285 | .003** | 0.02 | 0.10 |
| Non-autistic 2/5 | 0.10 | 0.02 | 4.38 | 1285 | <.001*** | 0.05 | 0.14 |
| Non-autistic 4/5 | 0.08 | 0.02 | 3.29 | 1285 | .001** | 0.03 | 0.12 |
| Non-autistic 4/3 | 0.04 | 0.02 | 2.35 | 1285 | .019* | 0.01 | 0.08 |
| Non-autistic 2 | -0.10 | 0.02 | -6.26 | 1285 | <.001*** | -0.13 | -0.07 |
| Non-autistic 3 | -0.10 | 0.02 | -5.90 | 1285 | <.001*** | -0.14 | -0.07 |
| Non-autistic 4 | -0.07 | 0.02 | -4.02 | 1285 | <.001*** | -0.11 | -0.04 |

Note. Model equation: DV ~ Condition + Group + (Condition x Group) + (1|Trial) + (1|Participant), 'Link', 'log'. Statistics as produced by the Matlab fitglm function. \*p < .05, \*\*p < .01, \*\*\*p < .001.

#### Autistic participants decompose shapes into a greater number of submovements at higher angular frequencies

For submovements there was no significant main effect of group (*F*(1,1285) = 3.50, *p* = .061). However, there was a significant main effect of condition (*F*(6, 1285) = 88.71, *p* < .001) and a group x condition interaction (*F*(6, 1285) = 6.31, *p* < .001). Coefficients from a mixed effects model (submovements ~ Condition + Group + (Condition x Group) + (1|Trial) + (1|Participant)), can be seen in **Supplementary Table 3**. The main effect of group indicates that autistic participants decomposed movements into a greater number of submovements compared to non-autistic participants. The main effect of condition indicates that, compared to the grand mean, low angular frequency shapes were decomposed into more submovements whereas higher angular frequency shapes were decomposed into fewer submovements. The *β* coefficients (**Supplementary Table 3**) and **Supplementary Fig. 1** illustrate a U-shaped relationship such that the fewest submovements were produced for the elliptical shape.

*β* coefficients relating to the interaction between group and condition illustrate that the mean estimated difference in submovements between groups varies as a function of condition. Notably, this interaction term was only statistically significant for conditions 4/5, 2 and 4. Thus, for all conditions aside from 4/5, 2 and 4, mean submovements can be relatively accurately predicted from the combination of the main effect of condition, plus an additional adjustment for group (e.g., intercept + autistic + 2/33). However, 4/5, 2 and 4 do not show this typical pattern. For 4/5, the difference between the groups is smaller than the difference that would be predicted from the main effects (non-autistic 4/5 is positive, also see **Supplementary Fig. 1**). For conditions 2 and 4 the difference between the groups is greater than the difference that would be predicted from the main effects (non-autistic 2 and non-autistic 4 are negative, see **Supplementary Fig. 1**)

**Supplementary Table 3.** Model Parameters for Submovements Mixed Models.

| | $\beta$ estimate | SE | t statistic | DoF | p value | Lower CI | Upper CI |
| --- | --- | --- | --- | --- | --- | --- | --- |
| **Submovements** |  |  |  |  |  |  |  |
| Intercept | 3.73 | 0.06 | 63.76 | 1285 | <.001*** | 3.61 | 3.84 |
| Non-autistic | -0.11 | 0.06 | -1.87 | 1285 | .061 | -0.22 | 0.01 |
| Autistic | 0.11 | 0.06 | 1.87 | 1285 | .061 | -0.01 | 0.22 |
| 2/33 | 0.50 | 0.03 | 15.91 | 1285 | <.001*** | 0.44 | 0.56 |
| 2/5 | 0.31 | 0.03 | 9.72 | 1285 | <.001*** | 0.24 | 0.37 |
| 4/5 | 0.11 | 0.03 | 3.51 | 1285 | <.001*** | 0.05 | 0.18 |
| 4/3 | -0.12 | 0.03 | -3.85 | 1285 | <.001*** | -0.19 | -0.06 |
| 2 | -0.45 | 0.03 | -13.88 | 1285 | <.001*** | -0.51 | -0.38 |
| 3 | -0.21 | 0.03 | -6.68 | 1285 | <.001*** | -0.28 | -0.15 |
| 4 | -0.14 | 0.03 | -4.18 | 1285 | <.001*** | -0.20 | -0.07 |
| Non-autistic 2/33 | -0.02 | 0.03 | -0.63 | 1285 | .528 | -0.08 | 0.04 |
| Non-autistic 2/5 | 0.05 | 0.03 | 1.59 | 1285 | .111 | -0.01 | 0.11 |
| Non-autistic 4/5 | 0.14 | 0.03 | 4.27 | 1285 | <.001*** | 0.07 | 0.20 |
| Non-autistic 4/3 | 0.06 | 0.03 | 1.82 | 1285 | .069 | 0.00 | 0.12 |
| Non-autistic 2 | -0.11 | 0.03 | -3.54 | 1285 | <.001*** | -0.18 | -0.05 |
| Non-autistic 3 | -0.03 | 0.03 | -0.84 | 1285 | .401 | -0.09 | 0.04 |
| Non-autistic 4 | -0.08 | 0.03 | -2.61 | 1285 | .009** | -0.15 | -0.02 |

Note. Model equation: DV ~ Condition + Group + (Condition x Group) + (1|Trial) + (1|Participant). Statistics as produced by the Matlab fitlme function. \*p < .05, \*\*p < .01, \*\*\*p < .001.

## Supplementary Material 3

### FFT of curvature confirms matched task compliance between groups

The FFT analysis was conducted in the curvature domain and revealed no significant differences between groups (see **Supplementary Fig. 2**). Given that this comparison of curvature values derived from the groups’ drawings identified no clusters of statistical significance, our findings indicate that task compliance was matched between autistic and non-autistic individuals i.e., compared to non-autistic participants, those with autism did not draw more/less curvy trajectories.

**Supplementary Figure 2.**
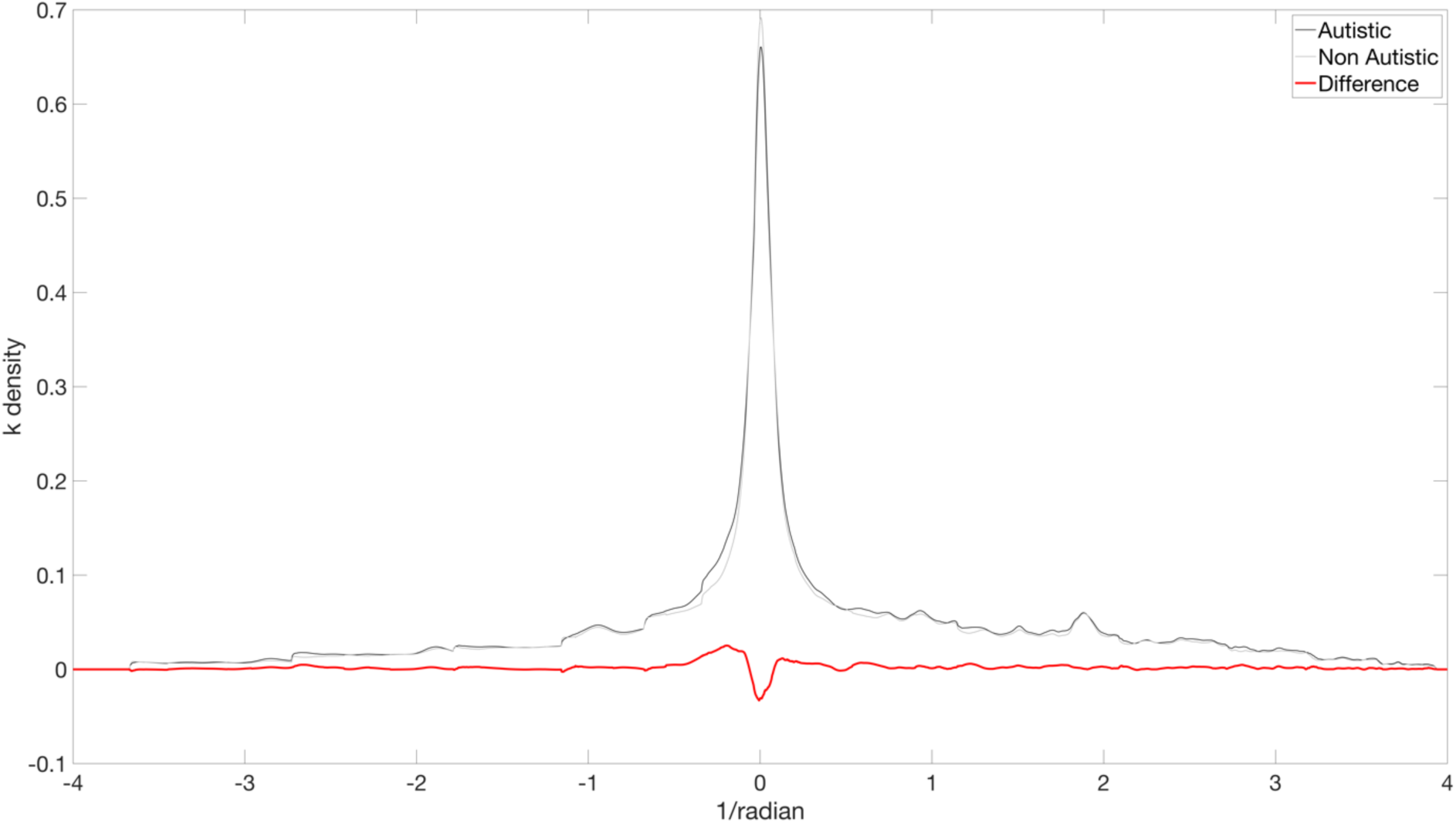
Amplitude Spectral Density Graph of Curvature. Amplitude spectral density of curvature for all angular frequency-defined conditions aligned to zero. Black line = autistic, Grey line = non-autistic, Red line = autistic - non-autistic.

## Footnotes

[1] Condition-first terminology is used throughout in line with the majority preference expressed surveys of the autistic community

[2] Also known as the two-third power law relating angular speed and curvature

[3] The angular frequency is defined as the number of oscillations per cycle. An ellipse has an angular frequency of two because in every 360° (or 2 pi radians) cycle of θ there are two points where the curvature of the trajectory changes dramatically (i.e., two “corners”). A square has an angular frequency of four corresponding to four changes in curvature in every 360° cycle. A logarithmic spiral has an angular frequency approaching zero because in every 360° cycle there is only a small (near zero) change in trajectory curvature.

